# rhizoTrak: A flexible open source Fiji plugin for user-friendly manual annotation of time-series images from minirhizotrons

**DOI:** 10.1101/547604

**Authors:** Birgit Möller, Hongmei Chen, Tino Schmidt, Axel Zieschank, Roman Patzak, Manfred Türke, Alexandra Weigelt, Stefan Posch

**Author notes:** These authors contributed equally (Birgit Möller, Hongmei Chen).

## Abstract

**Background and aims:** Minirhizotrons are commonly used to study root turnover which is essential for understanding ecosystem carbon and nutrient cycling. Yet, extracting data from minirhizotron images requires intensive annotation effort. Existing annotation tools often lack flexibility and provide only a subset of the required functionality. To facilitate efficient root annotation in minirhizotrons, we present the user-friendly open source tool rhizoTrak.

**Methods and results:** rhizoTrak builds on TrakEM2 and is publically available as Fiji plugin. It uses treelines to represent branching structures in roots and assigns customizable status labels per root segment. rhizoTrak offers configuration options for visualization and various functions for root annotation mostly accessible via keyboard shortcuts. rhizoTrak allows time-series data import and particularly supports easy handling and annotation of time series images. This is facilitated via explicit temporal links (connectors) between roots which are automatically generated when copying annotations from one image to the next. rhizoTrak includes automatic consistency checks and guided procedures for resolving conflicts. It facilitates easy data exchange with other software by supporting open data formats.

**Conclusions:** rhizoTrak covers the full range of functions required for user-friendly and efficient annotation of time-series images. Its flexibility and open source nature will foster efficient data acquisition procedures in root studies using minirhizotrons.

## 1 Introduction

Roots serve vital functions in the life of terrestrial plants. These functions include the provision of anchorage for the plant body, the uptake of nutrients and water from the soil (which in most cases is associated with symbiotic mycorrhizal fungi or bacteria), the transport of absorbed resources to the stem, and the storage of resources (Gregory, 2008). These functions are inherently associated with growth and death of roots and consequently linked to nutrient and carbon cycling in terrestrial ecosystems (de Kroon and Visser, 2003). Root biomass represents a substantial fraction of plant biomass, ranging from 16% in tropical forests to 77% in grasslands (Poorter et al., 2012). Root turnover accounts for approximately one third of the global primary productivity (Gill and Jackson, 2000). However, despite the importance of roots for plant growth and ecosystem functioning, they are less well studied than shoots due to their relatively lower accessibility.

This is especially true when it comes to studying the development of roots *in situ* and over time to characterize root growth and turnover. Multiple methods have been used to approach root turnover including ingrowth cores, sequential soil coring, nitrogen balancing, ^13^C labeling, and minirhizotrons (de Kroon and Visser, 2003; Lukac, 2012; Smit et al., 2000). However, sequential destructive coring is not always possible or desired and many ecological experiments request nondestructive approaches for root measurements such as minirhizotrons. Minirhizotrons are transparent tubes inserted in the soil through which roots growing at the interface between the tube and soil can be observed repeatedly using optical equipment (e.g., scopes, cameras, or scanners) (de Kroon and Visser, 2003; Smit et al., 2000). The major strength of the minirhizotron approach is that it enables repeated root measurements at the same position. This technique thus allows to distinguish between root production and mortality (Johnson et al., 2001; Majdi, 1996), which both occur simultaneously throughout the year (Hendrick and Pregitzer, 1992). Multiple parameters can be derived from time-series root images: standing root length, root diameter, root length production and mortality, and root life span.

However, extracting root data from minirhizotron images can be extremely time-consuming and laborious depending on the biological question. While for some questions it may suffice to measure overall root length of a given image, for more comprehensive investigations additional information like root diameter may be needed. The complexity of annotating minirhizotron images highly depends on the characteristics of the root images. In experimental settings, e.g., ecotrons, containers are filled with sieved, homogeneous soil creating images with high contrast between roots and background. Yet, in natural field conditions, the soil is usually heterogeneous often resulting in low contrast between roots and background. Also, a large number of roots with a high degree of overlap in an image adds to additional complexity. Finally, replicate minirhizotron tubes are recommended to account for the heterogeneous horizontal distribution of roots (Rewald and Ephrath, 2013). This exacerbates the already extensive effort for manual root annotation in root studies using minirhizotrons.

Over the years, numerous attempts have strived to ease the annotation task via automated image analysis (Erz et al., 2005; Zeng et al., 2010). However, only relatively simple scenarios, such as plants growing in agar plates and washed roots, are well tackled so far (Colombi et al., 2015; Nagel et al., 2012; Rellán-Álvarez et al., 2015). Tools for handling more complex minirhizotron images mostly concentrate on efficient and user-friendly functions for manual annotation by mouse point-and-click. Prominent examples of such tools include commercial software such as WinRHIZO Tron^1^ and RootSnap!^2^, and open source software, e.g., RootNav (Pound et al., 2013), DART (Le Bot et al., 2010), RootFly (Zeng et al., 2010), and SmartRoot (Lobet et al., 2011). All of them support a common set of basic editing operations but differ in their flexibility to represent branching roots, ways of handling time-series images, and the availability of advanced functionality. In Table 1 we present a comprehensive overview of basic features (e.g., options for adding and manipulating annotations), the model used for representing roots, and of more advanced functions (e.g., time-series support and data import and export). Below we will discuss the most important features and relevant differences depicted in Table 1 in more detail.

**Tab. 1.** Overview of existing tools for annotating root images and their basic and advanced features in comparison to our new tool rhizoTrak. For further explanations and specific details refer to the text.

| Feature | RootNav | Dart | RootFly | SmartRoot | RootSnap<sup>1</sup> | WinRhizo Tron | rhizoTrak |
| --- | --- | --- | --- | --- | --- | --- | --- |
| OS | Win | all | Win | all | Win<sup>m</sup> | Win | all |
| Terms of Usage | free | free | free | free | free<sup>n</sup> | comm | free |
| Language | C# .NET | Java | C++ | Java | n/a | n/a | Java |
| Source Code Available | (✓)<sup>a</sup> | — | ✓ | ✓ | — | — | ✓ |
| Root Tracing |  |  |  |  |  |  |  |
| Root Model | P<sup>c</sup> | T<sup>d</sup> | P | P<sup>c</sup> | T | T | T |
| (Semi-)automatic | ✓ | — | ✓ | ✓ | ✓ | — | — |
| Manual Annotation |  |  |  |  |  |  |  |
| Add/Delete Roots | (✓)<sup>b</sup> | — | ✓ | ✓ | ✓ | ✓ | ✓ |
| Move Nodes | ✓ | ✓ | ✓ | ✓ | ✓ | ✓ | ✓ |
| Merge/Split Roots | — | — | — | ✓ | ✓ | — | ✓ |
| Branching | ✓ | ✓ | — | ✓ | ✓ | ✓ | ✓ |
| Redirect Roots | ✓<sup>b</sup> | — | — | — | — | — | ✓ |
| Additional Data |  |  |  |  |  |  |  |
| Width | — | — | R | N | N | N | N |
| Status | — | — | R<sup>f</sup> | R, N<sup>h</sup> | R | N | N |
| Time Series Support |  |  |  |  |  |  |  |
| Load Series | — | — | ✓ | — | ✓ | — | ✓ |
| Copy Annotations | — | ✓<sup>e</sup> | ✓ | ✓<sup>i</sup> | ✓ | ✓<sup>i</sup> | ✓ |
| Temporal Links | — | — | ✓<sup>g</sup> | ✓<sup>j</sup> | n/a | ✓<sup>g</sup> | ✓ |
| Image Preprocessing |  |  |  |  |  |  |  |
| Enhancement | — | — | — | — | ✓ | ✓ | ✓ |
| Registration | — | — | — | ✓<sup>k</sup> | ✓<sup>k</sup> | — | ✓ |
| Data Exchange Formats |  |  |  |  |  |  |  |
| Import | — | csv | — | rsml | — | — | rsml |
| Export | txt,rsml,db | csv | csv | csv,rsml,db | csv | csv | csv,rsml |

Abbreviations used in Table 1

| Section | Name | Definition |
| --- | --- | --- |
| OS | Win | Windows |
|  | all | Win/MacOS/Linux |
| Terms of Usage | comm | commercial |
| Root Tracing | P | polyline |
|  | T | treeline |
| Additional Data | R | complete roots only |
|  | N | individual nodes |
| Data Formats | csv | comma separated values |
|  | txt | text file |
|  | rsml | root system markup language |
|  | db | data base |

Roots are often composed of different branches. For simplification, we refer to a single root branch as ‘root’ and denote a collection of connected branches as ‘root system’. In many tools, roots are represented by polylines composed of a set of nodes connected by linear segments. Each node in a polyline has maximal one incoming and one outgoing segment (Fig. 1). Hence, the topology of a real root system with branches cannot be represented directly with one polyline. SmartRoot models a root system with a set of polylines. Lateral roots can be attached to their parent root, however, this is not visualized in the graphical user interface. Very few tools, e.g., WinRHIZO Tron, use treelines to represent the topology of root systems. Similar to polylines, a treeline is comprised of a set of nodes linked by linear segments, but nodes in a treeline may be connected to more than one outgoing segment. Thus, treelines allow to directly represent root systems (Fig. 1). In addition to representing the topology of a root system, it is often required to annotate root segments with a status, e.g., a biological status like ‘alive’ and ‘dead’ (Fig. 1, small squares with letters). While WinRHIZO Tron supports to assign predefined status labels to each segment, other tools only allow for assigning a label to a complete polyline or treeline. This significantly reduces the accuracy in subsequent evaluations.

**Fig. 1.**
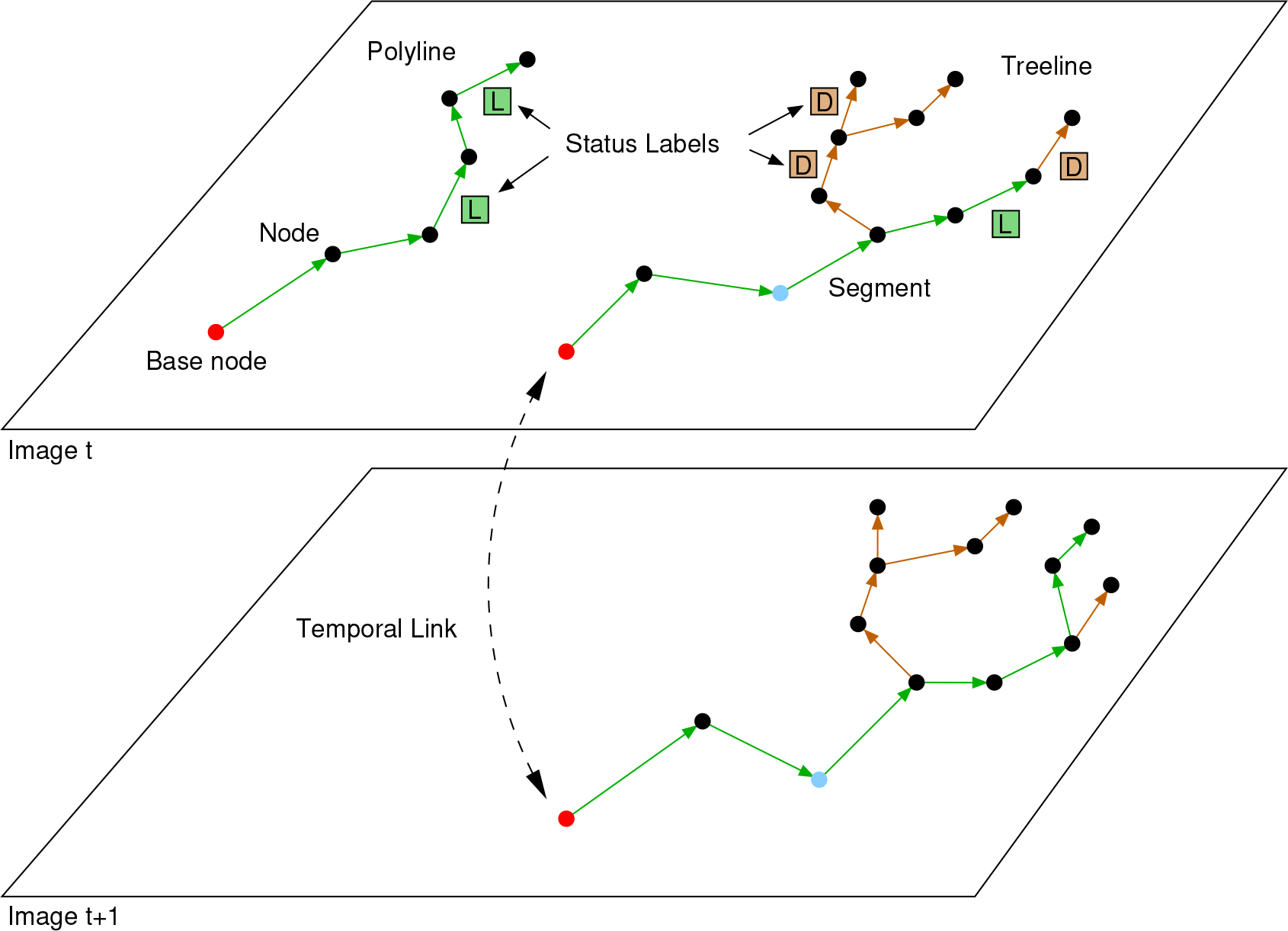
Schematic illustration of terms used in this article. Given are two images of a time series referring to time points t (top) and t+1 (bottom). The top image shows an annotation by polyline on the left and a treeline annotation on the right. Nodes are shown as black dots, the base nodes of each poly- or treeline are highlighted in red. The blue dot visualizes the changed position of individual nodes over time (see text for details). Arrows between nodes are referred to as segments. The status of individual segments is indicated by color, here green for living root segments (L) and brown for dead root segments (D), and by the status labels as colored squares next to the segments. Between both images the treelines are connected by a temporal link (double-headed dashed arrow) to represent that they belong to the same physical root. Note that temporal links are always visualized as emerging from the base node of a treeline. For further explanations, refer to the text.

One key advantage of the minirhizotron technique is the possibility to monitor individual roots over time. A proper support of time-series annotation is therefore crucial to maximize output from minirhizotron experiments. Time-series annotation should enable a user-friendly inspection of root changes between images and should support an easy addition of temporal links between projections of individual roots at different time points (Fig. 1). In addition, modifying annotations in a previous time point, e.g., to correct the topological structure, may be necessary and may require modifications also in other time points. However, time-series support is often limited in root annotation software.

In DART, each root system is represented by one single treeline which can only grow over time by adding new nodes and segments. The location of existing nodes can be changed, but such changes affect all time points automatically. For the treelines in Fig. 1 this would imply that all corresponding nodes in different time points (e.g., the two nodes marked in blue) always share the same position. Hence, DART ignores the possibility that the location of individual roots may shift over time, e.g., due to seasonal changes in edaphic conditions. WinRHIZO Tron allows for importing multiple images of a time-series at once where annotation data can be transferred from one layer to another via pattern files as an initialization. Temporal links between roots are established via automatically generated root identifiers. However, as images of a time-series are shown side by side, direct visual inspection of temporal changes is hampered. In addition, modifying and adding additional annotations in earlier time points with consistent temporal links is cumbersome.

The two most elaborate open source tools for minirhizotron time-series annotation are currently SmartRoot and RootFly. In SmartRoot, root annotations from the last time point can be imported via a previously saved file where treelines representing the same physical root at different time points are linked by root identifiers. Yet, these identifiers need to be individually specified by the user, which easily leads to errors and is only feasible for a small number of roots. In addition, SmartRoot only allows for the import of single images at a time which hinders visual inspection of root changes over longer time-series. In RootFly, a complete time-series of images may be imported into a single project allowing to easily switch between images of the time-series and to directly copy annotated roots to other time points. The root identifiers used for temporal links are generated automatically, however, cannot be modified by the user. This avoids mistakes but does not allow for straightforward changes of temporal links after propagating annotations from one time point to the next.

Ideally, software tools to annotate minirhizotron images are required to support an adequate representation of root systems by treelines and provide a flexible handling of temporal links including the option to modify annotations at previous time points. The tools should be characterized by a high degree of user-friendliness by being able to (1) load all images of a time-series at once to ease direct comparison of different images, (2) grant intuitive access to frequently used annotation and editing operations, and (3) support the annotation process by a user-friendly presentation in the graphical user interface. For many research questions it is also inevitable to (4) assign individual status labels to different parts of a root or root system. In addition, (5) flexible import and export of annotation data and measurements in common and open formats is preferable, particularly for data exchange with other software. So far, available tools only cover subsets of these features and, thus, leave room for further improvements.

In this paper we want to close this gap.

To increase the efficiency of manually annotating complex minirhizotron time-series images, we present the new tool rhizoTrak. It covers the full range of functionality required for flexible and user-friendly timeseries annotation. Currently no other available annotation tool can offer such a broad diversity and flexibility for manual annotation of complex time-series data.

rhizoTrak builds significantly on TrakEM2 (Cardona et al., 2012) and is realized as a Fiji plugin (Schindelin et al., 2012) which is available for Windows, Linux, and Mac OS. It can be easily installed using Fiji’s update site mechanism. rhizoTrak strictly follows open source paradigms and is released under GPL license, the source code is available from https://github.com/prbio-hub/rhizoTrak.

In the remainder of this article we present rhizoTrak’s main features and software architecture. First, we focus on basic design concepts, models, data structures, and implementation details of TrakEM2 and rhizoTrak, respectively (Materials & Methods). Second, we put a stronger emphasis on using rhizoTrak in practice and characterize the main features improving usability and annotation efficiency (Results). Finally, we summarize the main advantages of rhizoTrak over common software tools for root annotation available today, and outline rhizoTrak’s potential to foster efficient data acquisition procedures in root studies using minirhizotron images (Discussion).

## 2 Materials and Methods

The software basis of rhizoTrak is the open source software TrakEM2 (Cardona et al., 2012). TrakEM2 is dedicated to the semi-automatic reconstruction of neural circuits and combines basic image processing and analysis capabilities with manifold functionality for manual image annotation. On an abstract level, a root system shares important features with the dendrites of a neuronal cell. Both roots and dendrites are thin elongated and branching structures arising from the stem base and the cell body, respectively, and both do not form cycles. Rather, each root or dendrite has a unique direction pointing from the root base or the cell body, respectively, towards the tips of roots or dendrites. These topological similarities lead to comparable requirements for image annotation software in both domains and render TrakEM2 a viable foundation to develop an image annotation software for minirhizotron root images. Below we will outline the basic concepts and implementation details of rhizoTrak focusing on advanced features and extensions compared to TrakEM2.

### 2.1 Graphical user interface and time-series data

rhizoTrak aims to enable efficient and intutive annotation of complex time-series image data. Since manual image annotation is the main application of TrakEM2 it already provides manifold functionality for exactly this task. For example, TrakEM2 allows to import time-series data, offers a large collection of functions for adding and manipulating annotations, and provides options for easy data exchange with other tools. In addition, supplemental functions, such as enhancing image contrast and registering images, are provided. TrakEM2 offers a graphical user interface (GUI) to allow for direct access to the complete functionality.

rhizoTrak adopts the GUI of TrakEM2. Most of the time only a fraction of TrakEM2’s functionality is required for annotating roots in minirhizotron images. Since one of rhizoTrak’s main goals is a userfriendly annotation process, rhizoTrak additionally offers a lean mode of the GUI where only relevant functions and GUI elements are visible (Fig. 2). The lean mode improves usability and helps the user to stay focused on the root annotation task. The user can decide between the original TrakEM2 GUI and the lean interface via configuration options. The choice is stored in rhizoTrak’s project configuration file and will take effect when the user restarts the software.

**Fig. 2.**
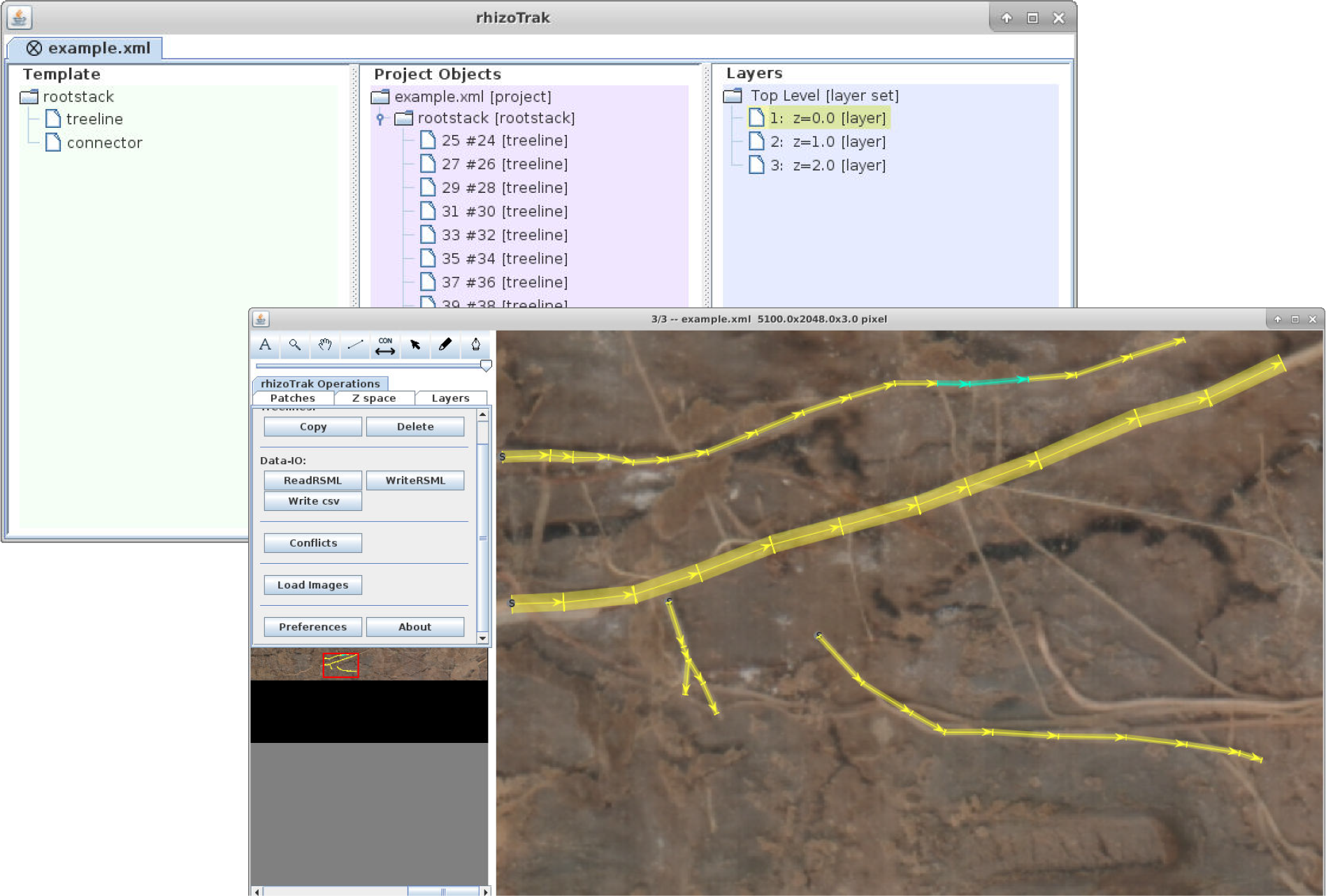
Screenshot of the rhizoTrak lean GUI with the project window (top left) and the layer window below containing a panel with manifold functionality on the left and the annotation work pane on the right (bottom). In the project window the overall project structure is depicted, in particular the different image layers are listed in the ‘Layers’ panel on the right and all annotation objects in the ‘Project objects’ panel in the middle. In the work pane a minirhizotron image with some annotated root systems indicated by the yellow and cyan treelines is displayed. The yellow and cyan color encodings of individual treeline segments refer to different status labels. Each node of a treeline can be assigned an individual diameter as indicated by the line segments passing through the nodes perpendicular to the segment orientation. The width of the treeline segments is interpolated from the diameters of their nodes and here visualized as colored quadrangles.

In neural circuit analysis 3D stacks comprising several image layers often need to be annotated. TrakEM2 allows to import all image layers of a stack into a single project and, thus, provides direct access to all layers during the annotation process. In a similar way, a time-series of minirhizotron images can be interpreted as a stack of image layers where each layer is associated with a time point. Hence, in rhizoTrak the images of a time-series are handled as one single project. 3D stacks are usually stored in a single file, while the different images of a time-series are saved in separate files with the file names encoding membership and temporal position within a series. rhizoTrak supports to import all files of a series at once. During import, the images can be sorted, either manually or automatically by acquisition date if they obey the ICAP naming convention (as introduced by the minirhizotron software of the Bartz Technology Corporation^3^, California, USA, see the rhizoTrak manual (available from the rhizoTrak website^4^) for details). Since minirhizotron experiments usually span a period of several months to years and new images become available gradually, rhizoTrak also supports the import of additional images into an existing project and enables an incremental annotation process. A search function allows to locate new images with ICAP file names that match the file names of the current project and presents them to the user for import.

### 2.2 Root annotation: treelines, status labels, and split and merge operations

Minirhizotron images often contain branching roots. To allow for straightforward annotation and direct representation not only of non-branching roots, but also of such complex root systems, rhizoTrak adopts treelines from TrakEM2. A treeline consists of nodes and segments (cf. Fig. 1). Each node can be augmented with an individual width to annotate root diameter.

Roots or a part of a root system in a minirhizotron image, i.e., the segments of a treeline, are often associated with a specific status, e.g., a biological status like ‘alive’ or ‘dead’. Such status information may be used, e.g., to assess root development over time by considering temporal changes in root status (Hendrick and Pregitzer, 1992) or to differentiate between roots originating from different species when they have distinct root morphology (Rewald et al., 2012).

rhizoTrak aims to offer the user a maximum of flexibility when dealing with status labels. In particular, each individual segment of a treeline can be assigned a separate status label, and the set of available status labels is fully configurable by the user. Internally a label is represented by a unique numerical identifier. Each label is associated to a name and abbreviation displayed as status label for selected treelines in the layer window of the GUI. By default, rhizoTrak defines the custom status labels LIVING (L), DEAD (D), DECAYED (Y), and GAP (G) which are often useful in minirhizotron image annotation. In addition, four more labels UNDEFINED, VIRTUAL, VIRTUAL RSML and CONNECTOR are defined for internal use. While these specific labels are fixed and cannot be modified, custom status labels including names and abbreviations can be customized by the user according to the biological question at hand. Custom status label definitions are stored in the project configuration and can be easily transferred to other projects.

All editing operations on nodes and segments (e.g., adding, deleting, or moving) as well as changes in segment status can be performed via mouse point-and-click and wheel actions, or via keyboard shortcuts.

rhizoTrak provides manifold functionality for modifying annotations and particularly for altering annotated root system topology consistently over complete time-series in an intuitive and efficient way. Two special features of rhizoTrak in this regard and rarely found in other tools are the support for altering the base node of a treeline and for treeline merges and splits. The base node of a treeline can be altered by declaring an arbitrary node of the treeline as new base node. This induces automatic redirections of all treeline segments on the path between the old and the new base nodes to recover a consistent root topology. Such alterations are particularly helpful in cases where it is difficult to correctly determine the growing direction of a root and the initial annotation might have been wrong.

Two treelines can be merged into a single treeline by adding a new segment between any nodes of the two treelines. The user needs to select the nodes via mouse. The order in which the nodes are selected defines the direction of the new segment and consequently the resulting topology of the merged treeline. The treeline to which the first selected node belongs is not modified and the second treeline is attached to the first treeline downstream of the selected node. If the second selected node is not the base node of that treeline, all segments connecting the base node and the selected node will be reversed for topological consistency after the merging (Fig. 3 a,b).

**Fig. 3.**
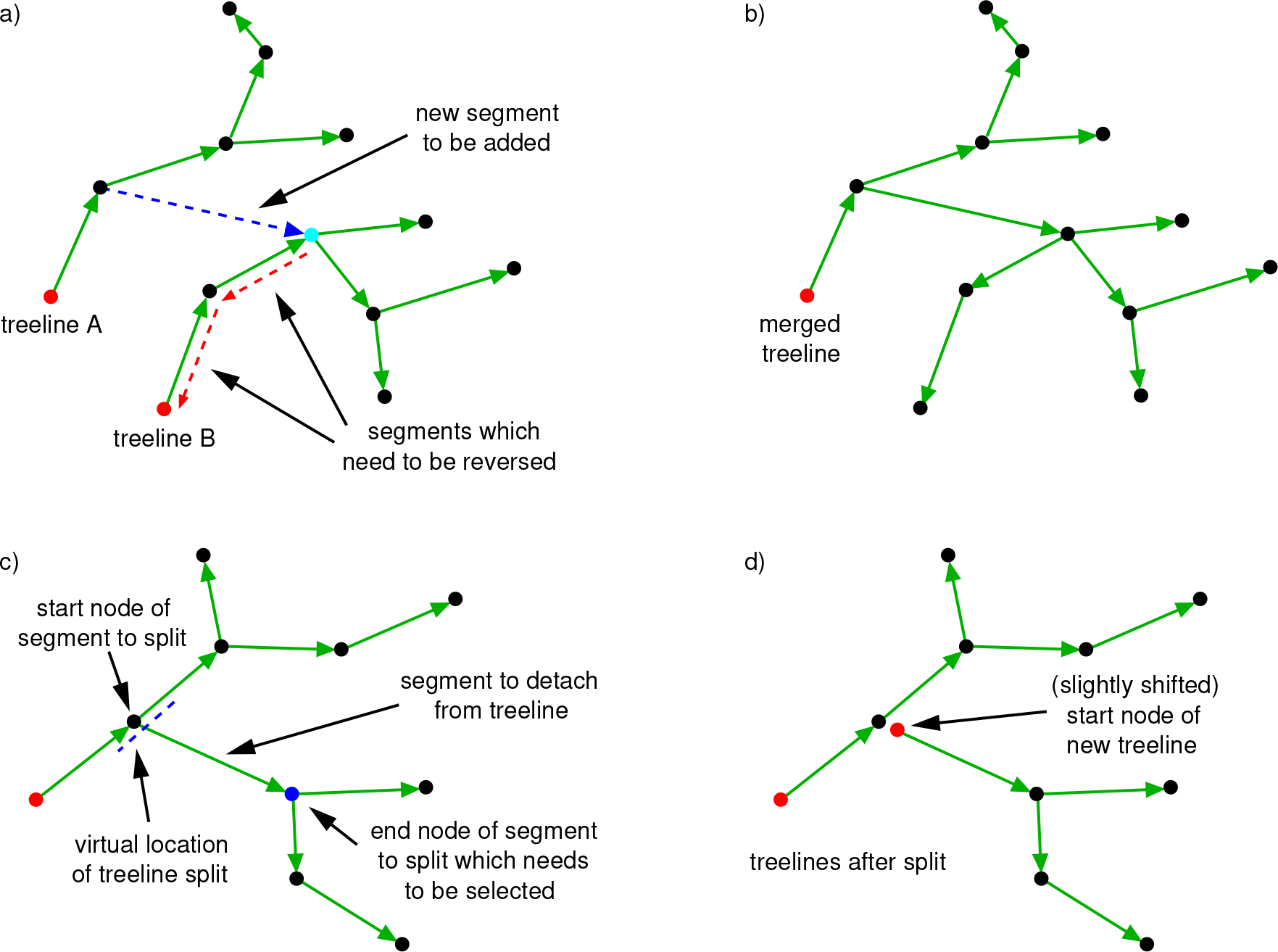
Merge and split operations on treelines. At the top, two treelines A and B are merged. The dashed blue segment indicates the new segment which links treeline B to treeline A (a). The merge requires all segments in treeline B connecting the former base node of B (shown in red) with the node where the link to treeline A is established (node marked in cyan, segments marked by dashed red arrows) to be reversed to recover a valid topology as shown top right (b). At the bottom, the treeline on the left is split up into two parts at the indicated segment ending at the blue node (c). The former start node of that segment is shifted along the segment direction to a new location (d) to avoid node overlap.

For splitting a treeline into two independent parts the user is required to select the end node of the segment which is to be detached from the treeline (Fig. 3 c, d). This will generate a new start node for the segment. This new start node will be placed in the vicinity of the old start node, however, slightly shifted along the direction of the segment to avoid overlap. The amount of shifting is determined by the maximum of the radius of the old start node and a pre-defined minimum distance between two nodes in case the radius is too small (Fig. 3 d).

### 2.3 Connectors and topological conflicts

A key question in minirhizotron image analysis is to distinguish between root production and mortality. This requires the distinction between different physical roots and to link annotations of treelines of the same physical root in images of different time points. rhizoTrak relies on a direct and fail-safe representation of such temporal relationships. The fundament of this representation is formed by the concept of *connectors* in TrakEM2. However, since TrakEM2 connectors are vulnerable to accidental manipulations by the user, rhizoTrak extends TrakEM2 connectors towards a larger flexiblity and reliability.

In TrakEM2, a connector is defined as a set of 2D positions over all image layers of a project and internally represented as a treeline in 3D space with each node of the connector treeline referring to one 2D position. The third dimension of each node is given by the index of the corresponding layer. Around each node of the connector, a circular 2D region is defined. Each annotation treeline overlapping with one of these circular regions becomes member of the group of treelines associated with the connector. Treelines can be added to or removed from such a group of treelines by altering the 2D positions of the connector nodes. However, since connector membership in TrakEM2 solely relies on geometric position, editing operations on treelines can easily result in accidentally adding or removing treelines to or from a connector. To overcome this problem, rhizoTrak relies on object identities rather than geometric locations to associate treelines with a connector. This extension decouples editing operations on treelines and the definition of connector membership.

If changes in root systems between successive images of a time-series are moderate, the annotation data from one image yields a reasonable initialization for the next. Hence, rhizoTrak allows to propagate annotations of one image layer to the subsequent layer. Moreover, in this case also connectors for corresponding treelines are automatically generated or extended. Over a complete time-series this may result in a significant reduction of manual annotation effort.

rhizoTrak represents each physical root in an image by one treeline, and all treelines in different images referring to the same physical root are grouped with a single connector. This implies two rules which have to be fulfilled for all connectors:

1. Each treeline may only be a member of one connector.
2. Each connector may not include more than one treeline from each image.

The connector concept also yields the basis for rhizoTrak’s flexible capabilities to alter root topology consistently over multiple time points. The two basic rules for connectors defined above allow rhizoTrak to automatically detect inconsistencies in root topology due to merges of treelines (e.g., Fig. 4, 5). rhizoTrak either automatically solves such topological conflicts (see Sec. 3.1 for an example) or asks the user to intervene. The integrated conflict manager lists all conflicts which need to be handled by the user. After a conflict is selected, the corresponding treelines and connectors are highlighted in the image and the user is guided through the resolution procedure.

**Fig. 4.**
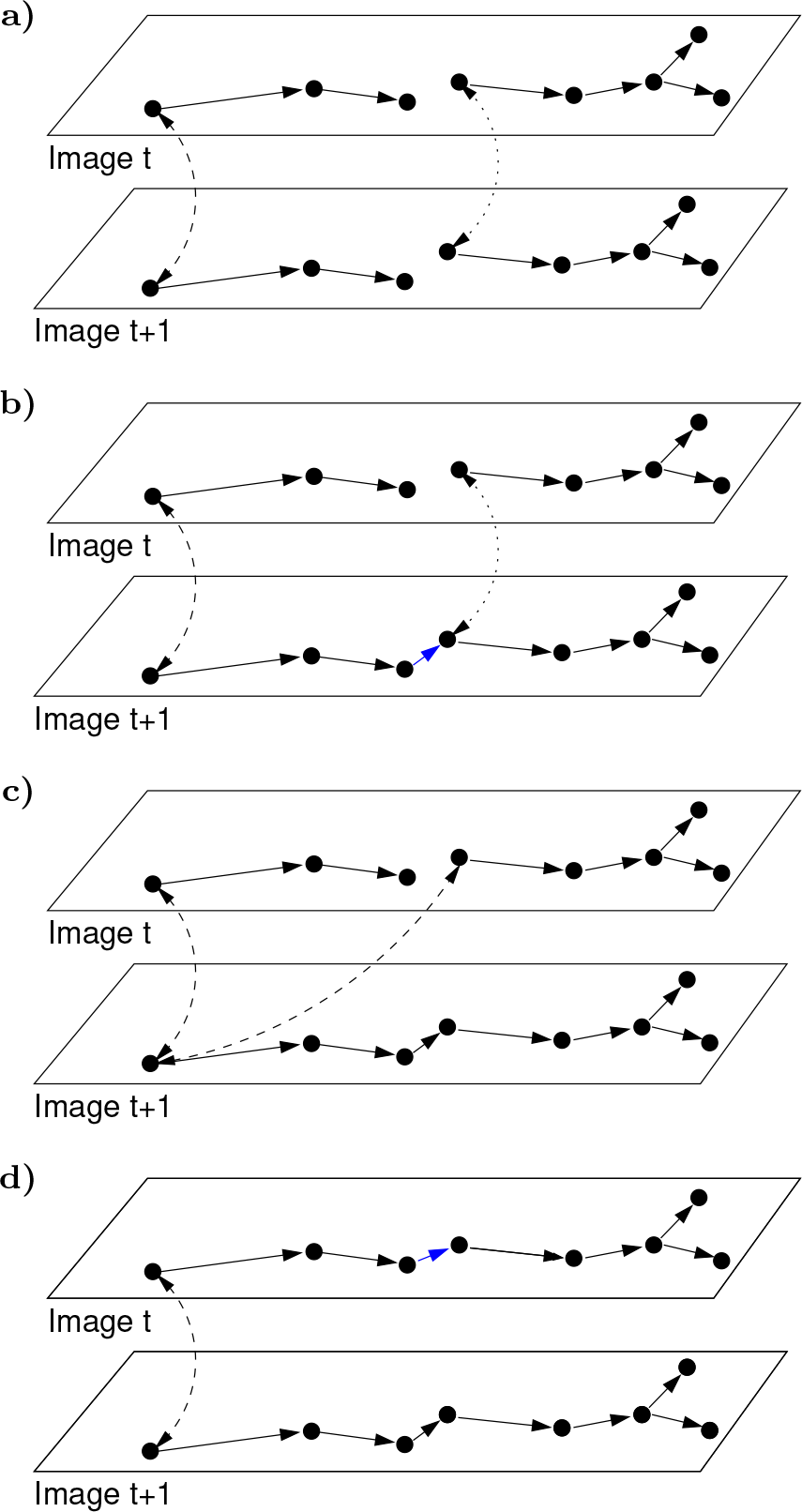
Example of a merge conflict and a possible solution. (a) In an image at time point t two root systems have been annotated. Then both annotations are copied to time point t+1 which triggers rhizoTrak to generate two different temporal links, i.e. connectors, between corresponding treelines shown as a dashed and dotted arcs. (b) The two treelines in image t+1 are then merged as indicated by the blue segment. (c) During the merging the two different connectors are also fused into a single one (as visualized by the dashed arcs) and the merged treeline of time point t+1 is linked to two independent treelines at time point t yielding a topological conflict. (d) One way to solve this conflict is to merge the treelines at time point t (as indicated by the new blue segment) which removes one node from the connector treeline and recovers topological consistency. For further details, refer to the text.

**Fig. 5.**
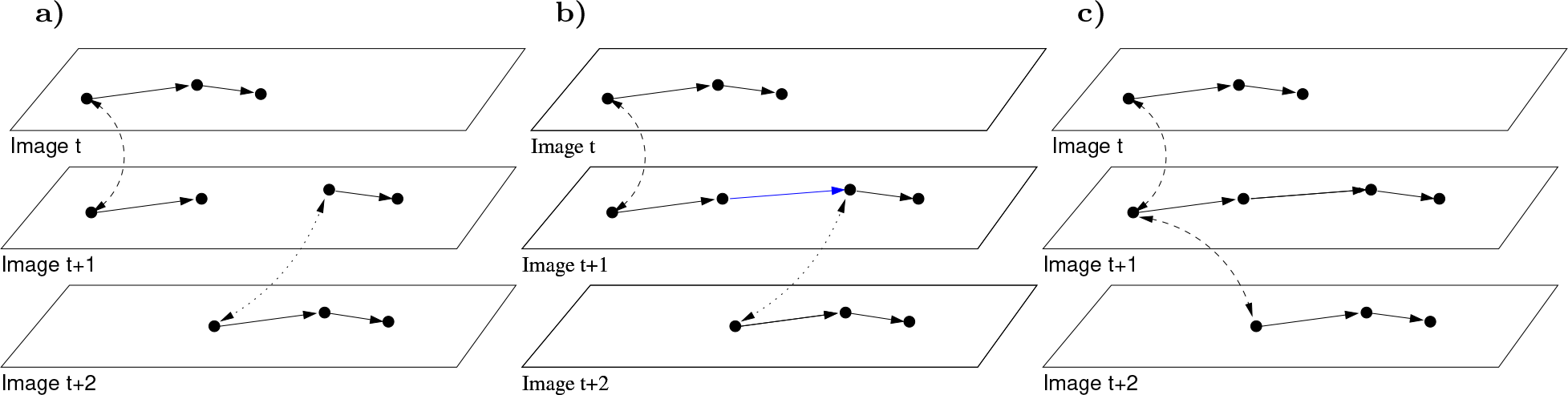
Merge conflict which can be automatically handled by rhizoTrak. Images of three time points are shown. (a) In the image of time point t, a single root is annotated, in the image of time point t+1 two roots are annotated, and in the image for the last time point again only one is annotated. In addition, two different connectors have been added as indicated by the dashed and dotted arcs. (b) If in time point t+1 the two roots are merged (as indicated by the blue segment) this results in a merged treeline with two different connectors being associated to that treeline, one linking to the root in image t and one linking to the root in image t+2. (c) By fusing both connectors into a single one (visualized by the dashed arcs) this conflict can be resolved. Each treeline is associated with a single connector again indicating that all three annotations are referring to the same physical root. See text for further explanations.

### 2.4 Root measurement and output

Research questions underlying the analysis of minirhizotron time-series data are manifold and root measurements of interest usually vary between different studies. rhizoTrak tackles these diverse requirements by supporting export of root measurements in a tab-separated format which can directly be used in further analysis, e.g., with common software tools like R or Excel. In the data files each row refers to an individual root segment (see Online Resource 1 for a sample data file). The first columns contain information about the experiment, tube identity, and time point to which an image refers, if such information can be extracted from the file names. Subsequent columns provide information about the segment itself, i.e., its ID, the ID of the root to which the segment belongs, the segment status, the length of the centerline, the diameters of the start and end nodes, and an estimate for the root surface area and volume. The ID of a segment is unique within each treeline and the IDs of treelines and connectors are unique within one project. Root IDs in the data files are either defined by the ID of the connector to which the treeline belongs or by the ID of the treeline in case a treeline is not member of a connector.

Root segments located completely outside of the valid image area are ignored when exporting root measurements. This might occur if annotations are propagated to a smaller image later in the timeline or if a treeline is shifted over the image boundary. If root segments intersect with the image boundary only the part inside is measured. Excluding parts of a treeline in measurements does not alter membership of the remaining segments to the treeline in data output.

For calculating geometric properties of a root segment like the surface area S and volume V, the root shape is assumed to form a truncated cone with radii *r*_*s*_ and *r*_*e*_ at the start and end nodes, respectively. The height of the cone h equals the length of the centerline. The segment volume V is calculated as:

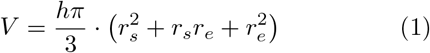

The segment surface area S equals the lateral area of the truncated cone and is calculated as:

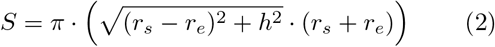

In addition to measurements of length, surface and volume for individual segments, aggregated measurements for all segments per time point and status label can be extracted (see Online Resource 2).

### 2.5 Annotation export in RSML

To facilitate easy data exchange with other tools rhizoTrak strictly follows an open access strategy adopting common data formats whenever reasonable. For import and export of root annotation data it supports the Root System Markup Language (RSML). RSML has been devised to yield a common basis to represent root annotation (Lobet et al., 2015). This facilitates annotation exchange between tools featuring RSML support, e.g., RSML adapters for Excel and R, for further processing and analysis. In RSML the annotation of each image along with its properties is stored in a separate RSML file. In this annotation the image is denoted as a scene, which is modelled to contain one or more plants and each plant is modelled to have one or more root systems (branching roots). A root system is represented as a hierarchically ordered set of non-branching roots. The geometry of a nonbranching root is represented with a polyline and may be augmented by so-called *functions* storing diameter or status labels. Additional properties or annotations may be added to each scene (image), plant, or root.

In rhizoTrak one or multiple RSML files can be imported in one single operation. An RSML file may be imported into a newly created layer or into an existing empty (first) layer. An RSML file can also be imported into a layer with an image already loaded. In this case rhizoTrak validates that this image and the one specified in the RSML file are identical using their hash codes where the SHA-256 codes are used as specified by RSML.

On import into rhizoTrak each root system in the RSML file is converted to one single treeline. As suggested by the RSML thesaurus (RSML Homepage, 2015, Section ‘Format’), rhizoTrak uses the parent-node property of an RSML root to define the connecting node in the parent root which has higher order in the hierarchy than the child root. If there is no parent-node property, the nearest node in the parent root to the base node of a child root is used as the connecting node. In addition to the polyline geometry, diameter and status label integers of nodes are imported if stored as RSML functions. Several examples on the RSML homepage (RSML Homepage, 2015, Section ‘Examples’) and shipped with the RSML conversion tools (RSML Github Repo, 2018, Project ‘RSML-conversion-tools’) represent the sample values of functions as text content of the corresponding HTML elements. This is in contrast to the Extensible Markup Language (XML) schema definition of RSML which uses attributes of the HTML elements instead. rhizoTrak supports both the text and the attribute form of the XML representation on import. An RSML file may define a mapping of status label integers to string representations in the property list of the scene. On import, rhizoTrak checks the consistency of the mappings defined in the current project and the RSML files. If inconsistencies with regard to images or status label mappings are detected, the user may cancel the import or proceed using the definitions already present in the project.

Since rhizoTrak features no concept of plants with multiple root systems, this information is not available to rhizoTrak users, however, preserved on subsequent re-export as required by the RSML specification. After importing all RSML files, rhizoTrak checks if all imported RSML files indicate that temporal links are represented. This is the case if all unified flags in the time series meta data are true in the RSML files. In this case connectors are established among treelines accordingly.

For exporting annotations into RSML files the active or all layers may be selected. Each layer is exported into one RSML file. The status label mapping of the current project is saved in the property list of the scene. If a treeline has been previously imported, its membership to a plant is preserved (see above). Each newly added treeline is exported as one newly created plant. A treeline is split into a set of non-branching polylines. If the branches have been created by user actions within rhizoTrak or have been read from an RSML file using the parent-node property, the hierarchic relationships between polylines are saved with RSML’s optional parent-node property. For each node the diameter and status label integer is exported in functions. If the geometry of a previously imported treeline has not changed after import, additional features like RSML functions, properties and annotations are re-saved as required by RSML specifications.

## 3 Results

### 3.1 Root annotation with rhizoTrak in practice

Fig. 2 depicts the graphical user interface of rhizoTrak which consists of two windows. The project window (Fig. 2, top left) keeps track of all annotation objects, and the layer window (Fig. 2, bottom) displays the images and all functionality to perform annotation tasks. Annotating a time-series of minirhizotron images starts with importing the images^5^. rhizoTrak supports to import the complete set of images of a time-series at once and handles a series of images as layers of a stack. The stack can be inspected in a comfortable and user-friendly way by scrolling through the different layers via mouse wheel. This allows for direct visual comparison of roots in different images, given that the different images of a minirhizotron time-series are properly registered, i.e., potential shifts and deformations between subsequent images have been corrected and all images refer to the same physical location. Moreover, the transparency of an image layer can be adjusted to enable overlay of different image layers.

Roots in an image can be annotated by mouse point-and-click actions or via keyboard shortcuts. rhizoTrak models roots as treelines and represents the topology of a root system with a single treeline object. rhizoTrak supports all common editing operations on treelines like moving, adding or deleting individual nodes. In addition, complete treelines can be repositioned, and status labels of subtrees can be changed with a single operation. One of the special features of rhizoTrak is the support for changing the base node of a treeline and, in particular, for split and merge operations on treelines. By a merge operation two formerly independent treelines in one image can be joined to form a single treeline. Vice versa, the split operation allows to split up a single treeline into two independent treelines.

rhizoTrak supports to augment each node of a treeline with a diameter and annotate each individual segment with a customizable status label. In addition, for each status options like display color or opacity can be configured by the user which are used for visualizing the segments. The diameter of a node is by default visualized as a line through the center of the node perpendicular to the centerline of the corresponding segment (Fig. 6), and the projected area of the root segment is visualized by a quadrangle formed by the two diameter lines and two lines connecting their end-points (Fig. 6). The user may choose to fill the quadrangle with the selected status color and opacity (Fig. 7 a) and/or choose to draw the outline (Fig. 7 b). The diameter can either be omitted or shown as line, circle or a combination of both (Fig. 7 a-c). Individual as well as all treelines can be set visible or invisible to ease root annotation in image areas where root density is high.

**Fig. 6.**
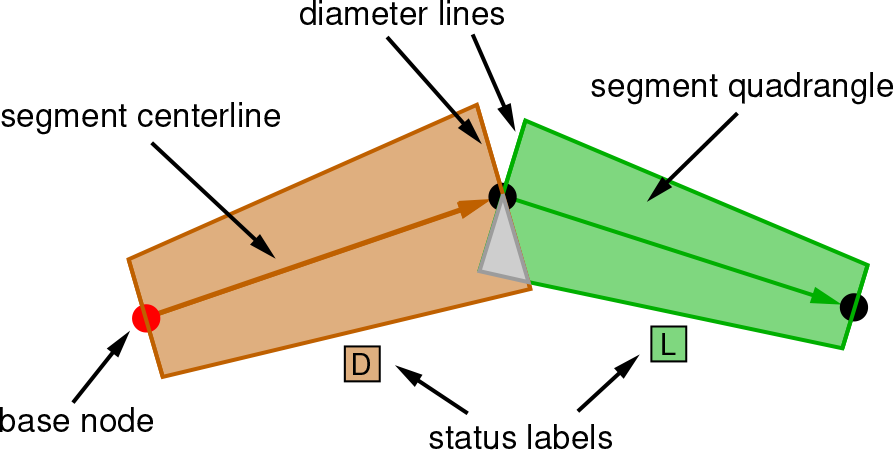
Root segments are by default visualized with their centerlines connecting the centers of their start and end nodes. The diameter of a node is indicated by a line through its center perpendicular to the segment centerline. The projected area of the root is shown as quadrangle formed by the diameter lines and lines connecting their end points. The color of each segment and the quadrangle area refer to the status of the segment, in this case green for living and brown for dead. For the currently selected treeline the status is also indicated by displaying the status label abbreviations close to the segment (L for living and D for dead). To ease comparison with underlying image data, segment visualization can be altered as shown in Fig. 7.

**Fig. 7.**
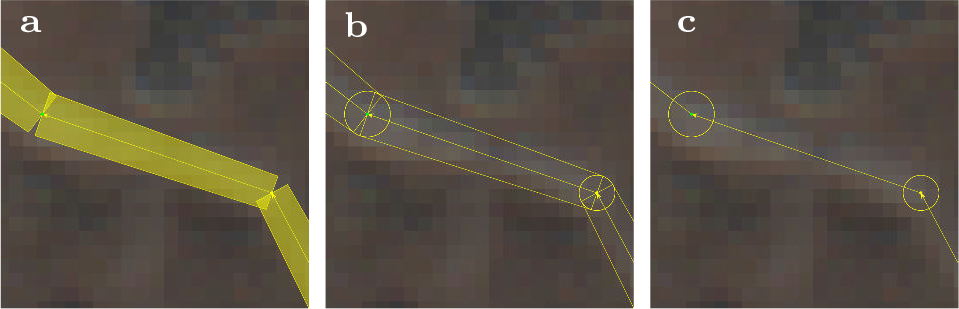
Different visualization options for segments in rhizoTrak. In a) and b) segment areas are visualized by quadrangles formed by the diameter lines of the nodes and line segments connecting their end-points. a) The quadrangle areas are filled with the color representing the segment status, b) just the quadrangle outlines and the nodes are visualized by their diameter lines and corresponding circles. In c) visualization is restricted to the node circles and segment centerlines to ease comparison of annotations with the underlying image.

Similar to other tools (e.g., RootFly, SmartRoot and WinRHIZO Tron), rhizoTrak allows to copy root annotations of one image layer to the next image layer as initial annotation. This automatically creates a temporal link (i.e., connector) between treelines which represent the same physical root in both layers. In contrast to other tools these connectors are independent of treeline identifiers which provides larger flexibility in further editing.

Merging treelines which are linked to a connector can result in topological inconsistencies. Consider for example that two treelines were annotated in the first layer of a stack and copied to the next layer (Fig. 4 a) which automatically creates two different connectors (dashed and dotted lines in Fig. 4 a). In the second layer the user decides to merge these treelines (blue segment in Fig. 4 b). As a result, the two different connectors are fused into a single connector (dashed arc in Fig. 4 c) for the merged treeline in the second layer which is linked to two different treelines in the first layer (Fig. 4 c). This reflects a topological inconsistency as a single physical root in one layer (time point t+1) cannot be identical to two different roots in another layer (time point t). The conflict may be solved by merging the treelines in the first layer as well (Fig. 4 d). rhizoTrak automatically monitors all merge operations and keeps track of such topological inconsistencies. Some of these inconsistencies can be resolved automatically (see Fig. 5 for an example) and rhizoTrak automatically propagates the changes in root topology to other time points. For situations where automated handling of a conflict is not possible (as in the example in Fig. 4) rhizoTrak features a conflict editor to inform the user about inconsistencies and guides possible solutions.

Root annotations can be imported and exported in RSML format (Lobet et al., 2015). One or multiple RSML files can be imported in a set of consecutive layers. Prior to the actual import, several consistency checks are performed (see Subsec. 2.5) and import may be cancelled if these checks fail. In addition to the geometry of roots, the diameters and status label integers of nodes along with status label mappings are imported if available in the RSML file. If no status labels are present in the RSML file, a user-defined default is used. Likewise, if information on temporal links between root systems at different time points is present, it is appropriately represented with connectors in rhizoTrak.

The current or all layers may be (re-)exported to RSML files. The user may choose if temporal links are exported. If an RSML file has been imported into a layer to be exported, additional optional properties, functions, and annotations previously read are retained as required by the RSML specifications. In addition, quantitative data from root annotations can be extracted to text files for further analysis with pertinent software tools.

### 3.2 Experimental evaluation

#### 3.2.1 Analyzing minirhizotron images using rhizoTrak and WinRHIZO Tron

To examine the reliability of rhizoTrak, we annotated two sets of example minirhizotron images^6^ using both rhizoTrak and WinRHIZO Tron MF (version 2014, Regent Instruments Inc. Québec, Canada) and used the output from WinRHIZO Tron as a reference. The first set of example minirhizotron images is a subset of a time series acquired with CI-600 In-Situ Root Imager (CID Bio-Science, Inc., Washington USA) at the Jena Experiment (Weisser et al., 2017). The original images were captured every other week from February to October in 2015. We selected five time points and cropped the images to the size of 6 × 3 cm for this analysis. The selected images showed the whole root life cycle (initiation, elongation, and disappearance) and demonstrated the changes and complexity of the background scene (Fig. 8 a,b). The second set of example images was acquired from an Ecotron experiment^7^ with the same scanner where soil was more homogeneous. This Ecotron experiment was designed to investigate the responses of ecosystem functions of grasslands to insect decline. We selected 20 minirhizotron image areas of the same size and resolution as the first set of example images, but varying in root length density (Fig. 8 c,d).

**Fig. 8.**
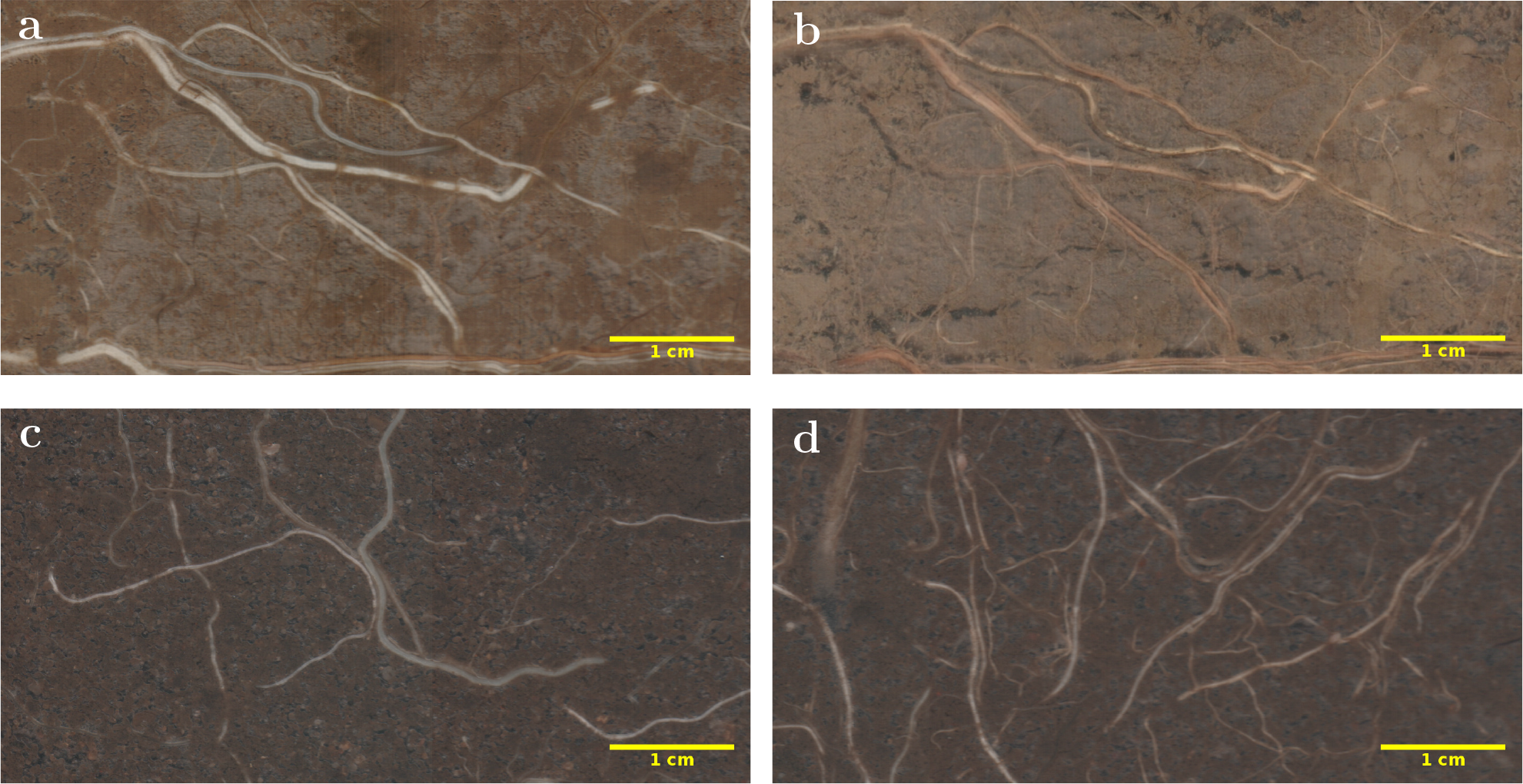
Examples of minirhizotron images for comparative analysis between WinRHIZO Tron and rhizoTrak. The images on top were taken in February (a) and June (b) in a grassland biodiversity experiment in Jena (Germany). The images at the bottom were acquired in an Ecotron experiment in Bad Lauchstädt (Germany) with low (c) and high (d) root density.

We pooled the measurements from these two sets of examples and examined the agreement between rhizoTrak and WinRHIZO Tron using the Passing-Bablok regression (Passing and Bablok, 1983) in the ‘mcr’ R package (Manuilova et al., 2018). The intercept was −0.09 cm cm^−2^ with a 95% confidence interval of [−0.24, 0.02]. The slope was 1.00 with a 95% confidence interval of [0.97, 1.06]. Thus, rhizoTrak and WinRHIZO Tron provided comparable measurements (see Fig. 9).

**Fig. 9.**
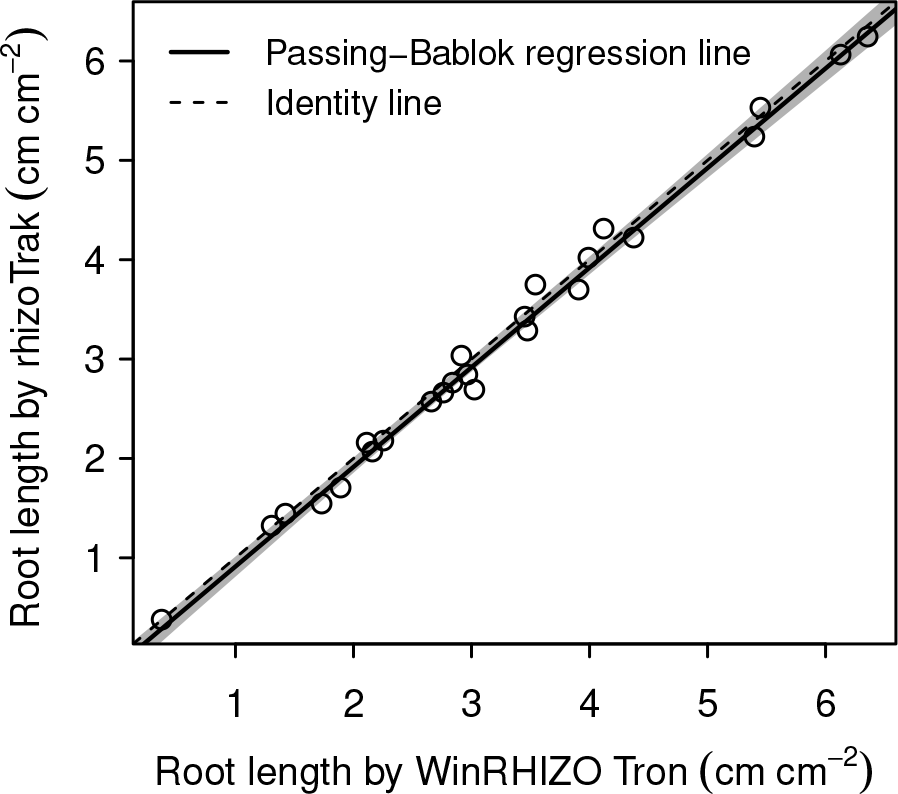
Passing-Bablok regression between root length measured by rhizoTrak (Y) and that measured by WinRHIZO Tron (X): Y = −0.09 + X. The grey zone represents the 95% confidence interval of the regression.

#### 3.2.2 Data exchange between rhizoTrak and other tools using RSML

The RSML format was developed to foster exchange between different annotation tools, image analysis tools, and modelling platforms (Lobet et al., 2015). To show rhizoTrak’s conformance with these ideas and the RSML standard, we imported two RSML examples (RSML Homepage, 2015, Section ‘Examples’) beetroot 0.rsml and beetroot 1.rsml, which belong to one time-series, into rhizoTrak. The geometry and diameters were correctly converted and the identity of roots in both images were represented by connectors. Subsequently the annotation of one treeline was modified, where nodes were displaced and additional branches were added. Then both layers were re-exported into two RSML files.

These RSML files generated by rhizoTrak were imported into SmartRoot (Lobet et al., 2011). This import again showed correct geometry and diameter of roots including the modifications performed within rhizoTrak.

Finally, the rhizoTrak annotations exported to RSML format were read by the R function (R Core Team, 2014) supplied as one of the RSML conversion tools (RSML Github Repo, 2018, Project ‘RSML-Validator’). Although the RSML files generated by rhizoTrak conformed to the RSML specifications and passed the schema validator RSMLValidator which is part of the RSML supplied conversion tools, reading the RSML file in R failed. This was caused by the status label mappings stored as scene properties. If these property definitions are removed from the RSML file, reading was successful and the annotations can be analyzed with the R functions.

## 4 Discussion

rhizoTrak is an open source tool which enables flexible and user-friendly manual root annotation of minirhizotron images. rhizoTrak is developed in Java on the basis of TrakEM2 (Cardona et al., 2012) and realized as a plugin for Fiji, which is compatible with all mainstream operating systems (Windows, Mac OS, and Linux) (Schindelin et al., 2012). rhizoTrak uses treelines instead of polylines to directly represent the branching structure in root topology. For treelines, diameter and user-defined status can be specified for each segment. rhizoTrak allows to import complete sets of time-series images which eases decisions on root status and structure by considering multiple images at the same time. In addition, rhizoTrak uses temporal links (connectors) to directly link treelines of the same physical root in different images over time rather than linking them indirectly via treeline identifiers as done in many other root annotation tools. The strict separation between treeline identifiers and temporal links enables the user to modify root topologies during annotation via merging or splitting treelines without losing temporal connections. A further useful function is the redirection of treelines since the upstream and downstream direction of a root is often not straightforward in minirhizotron images. All root measurements can be exported in a csv file and aggregated at the image level on data output.

rhizoTrak features two main advantages: (1) an elaborate support to annotate time-series images and (2) user-friendly and handy procedures for adding and manipulating root annotations. Minirhizotron images often confront the scientist with difficult choices due to the heterogeneity of the soil background and the variation of root growth and texture. Moreover, images are often analysed over one or two vegetation periods with naturally large differences in soil moisture and animal activity leading to severe changes in soil colour or air bubbles in the images. For much the same reason, identical roots often slightly shift position in addition to growing, senescing (often accompanied by changes in colour), shrinking or dying over time. Thus, it is paramount for a minirhizotron annotation tool to ease user decision making and enable maximal flexibility in the annotation process. Most often, considering multiple images of a time-series will help to make such decisions. In rhizoTrak, all images can be imported into a project at once. Thus, scrolling through the images (via mouse wheel or keyboard shortcuts), the user can verify the presence of a transparent newly born root, root growing direction, and root topology. In addition, overlaying images directly visualizes problems such as missing pixels or a few stretched pixels (due to brief hung-up of the scanner). Moreover, the user can copy the annotation of the current image to the next one and use it as a starting point for the annotation. After treeline propagation, the hide/unhide function (which can be easily toggled via a keyboard shortcut) provides an easy way to inspect the changes in roots and eliminates the necessity to drag the treeline away from the root below in the image for visual inspection. Elongating existing roots, changing segment diameter or status, altering the root position and many other editing operations are easily done via single keyboard operations, via mouse, or keyboard and mouse combined. These one-action operations will save a lot of time once the user gets familiar with rhizoTrak shortcuts.

By using connectors, rhizoTrak finds a balance between efficiency and flexibility for establishing and altering temporal links. Most available software tools represent temporal links with unique identifiers for treelines or polylines in different images, which is error-prone and renders changing earlier annotations tedious. For example, in WinRHIZO Tron, the identifier of a treeline changes from ‘RX’ (X represents a number) to ‘1’ when the base node of the treeline is deleted accidentally. Consequently, the temporal links for all treelines with the identifiers ‘RX’ and ‘1’ are compromised. In SmartRoot temporal links are specified manually assigning the same identifiers to different roots. However, specifying identifiers manually is tedious and prone to mistakes especially when the number of roots increases. In rhizoTrak, temporal links, i.e. connectors, are automatically generated or extended when treelines are copied from one image to the next. Subsequent changes at the treeline level do not affect the connectors in rhizoTrak. A treeline can be attached to or detached from a connector independent of its current position or structure. Furthermore, corrections of wrongly specified root topology in one image by merging treelines may result in topological inconsistencies in other images which will be listed in the conflict manager to assist the user to solve them. Thus, a comprehensive set of functions eases image annotation with rhizoTrak especially to correct previous decisions.

We tested rhizoTrak against WinRHIZO Tron, one of the most elaborate but commercial software tools available (see Table 1) and found no significant differences in root length measurement between both tools. From this and our experience in root annotation with both tools, we feel comfortable that rhizoTrak is a highly efficient and reliable tool to annotate minirhizotron images. Furthermore, due to import and export of annotations using the RSML specification, rhizoTrak features interoperability with other root software.

rhizoTrak offers an accessible and efficient way to annotate time-series minirhizotron images. Compared to existing tools which cover only a subset of the functions required for time-series annotation, rhizoTrak provides the entire spectrum of functionality essential for an intuitive, user-friendly, and time-efficient annotation process. rhizoTrak’s open source nature as a plugin for Fiji will largely simplify data acquisition in root studies using minirhizotrons in the future and enable the scientific community to develop useful add-on packages and provide additional features necessary for future research. Current work is focussed on further reducing the workload of annotating minirhizotron images by integrating automatic segmentation techniques into rhizoTrak. Already now rhizoTrak ships with an extensible interface definition for root segmentation algorithms which allows everybody to implement and directly integrate own segmentation approaches into rhizoTrak in a straightforward manner.

## Supporting information

Online Resource 1: sample measurements in detailed format

Online Resource 2: sample measurements in aggregated format

## Electronic Supplementary Material

The manuscript is supplemented with the following two online resources:

- Online Resource 1: ESM 1-rhizoTrak.csv This file contains sample root measurements in detailed CSV format. Each row contains data for an individual root segment.
- Online Resource 2: ESM 2-rhizoTrak.csv This file contains sample root measurements in aggregated CSV format. For each layer (from 1 to 5 in this case) aggregated measurements are given, grouped according to segment status (here LIVING, DEAD, DECAYED or GAP).

The files can also be downloaded from the ‘Sample data’ section on rhizoTrak’s homepage, https://prbio-hub.github.io/rhizoTrak/pages/data.html.

## Footnotes

a (Pound et al., 2013) state that the software is open source, but source code could not be found

c implicitly via manipulating control points

d polylines can be grouped hierarchically

e one single treeline per root system for a time series

f inherent due to model of one treeline per root system over all time-points

g only DEAD/LIVING

h via automatically generated IDs

i assignable to nodes and intervals

j via files

k manually adopting user-defined identifiers

l manually

m all information given here based upon the manual

n requires a touch device

o registration required

1 WinRHIZO Tron website, accessed 15 Jan 2019, http://regent.qc.ca/assets/winrhizo_about.html

2 RootSnap! website, https://cid-inc.com/plant-science-tools/root-measurement-with-minirhizotron/ci-600-in-situ-root-imager/#rootsnap, accessed 15 Jan 2019

3 Bartz Technology Corporation, ICAP software, https://www.vienna-scientific.com/products/minirhizotron-systems/mr-software-icap/, accessed 15 Jan 2019

4 https://prbio-hub.github.io/rhizoTrak/

5 A sample rhizoTrak project including a small time-series of minirhizotron images can be found on rhizoTrak’s website, https://prbio-hub.github.io/rhizoTrak/.

6 Sample images can be found at rhizoTrak’s webpage, https://prbio-hub.github.io/rhizoTrak/.

7 The iDiv Ecotron, https://www.idiv.de/en/research/platforms_and_networks/idiv_ecotron.html, accessed 15 Jan 2019

